# Novel 2D/3D Hybrid Organoid System for High-Throughput Drug Screening in iPSC Cardiomyocytes

**DOI:** 10.1101/2024.04.29.591754

**Authors:** Jordann Lewis, Basil Yaseen, Anita Saraf

**Author notes:** **Corresponding Author:** Anita Saraf, MD, PhD, John G. Rangos Sr. Research Center, #8122, 4401 Penn Ave, Pittsburgh, PA 15224.

## Abstract

Human induced pluripotent stem cell cardiomyocytes (hiPSC-CMs) allow for high-throughput evaluation of cardiomyocyte (CM) physiology in health and disease. While multimodality testing provides a large breadth of information related to electrophysiology, contractility, and intracellular signaling in small populations of iPSC-CMs, current technologies for analyzing these parameters are expensive and resource-intensive. We sought to design a 2D/3D hybrid organoid system and harness optical imaging techniques to assess electromechanical properties, calcium dynamics, and signal propagation across CMs in a high-throughput manner. We validated our methods using a doxorubicin-based system, as the drug has well-characterized cardiotoxic, pro-arrhythmic effects. hiPSCs were differentiated into CMs, assembled into organoids, and thereafter treated with doxorubicin. The organoids were then replated to form a hybrid 2D/3D iPSC-CM construct where the 3D cardiac organoids acted as the source of electromechanical activity which propagated outwards into a 2D iPSC-CM sheet. The organoid recapitulated cardiac structure and connectivity, while 2D CMs facilitated analysis at an individual cellular level which recreated numerous doxorubicin-induced electrophysiologic and propagation abnormalities. Thus, we have developed a novel 2D/3D hybrid organoid model that employs an integrated optical analysis platform to provide a reliable high-throughput method for studying cardiotoxicity, providing valuable data on calcium, contractility, and signal propagation.

## Introduction

Despite rigorous preclinical and clinical pharmaceutical testing, a majority of drugs do not meet relevant endpoints to translate to market.^1^ Among drugs that are proven safe and effective, many eventually get removed from the market due to off-target toxicity.^2,3^ Of particular interest is cardiotoxicity, as many drugs can affect the heart, inducing heart failure, cardiomyopathy, arrhythmias, and other cardiac pathologies.^3-5^ Thus, there is a need for an improved, high-throughput drug screening method which can distinguish effects on cardiac tissue. In recent decades, cardiomyocytes differentiated from human induced pluripotent stem cells (hiPSC-CMs) have provided a platform to study various cardiovascular disease pathophysiology, as well as drug efficacy and toxicity.^6-8^ While 2D cell culture offers a rapid and relatively simple model to study CM structure and function, it does not completely emulate cardiac tissue, with low complexity and not recapitulating the multitude of cell-cell interactions.^9^ In recent years, three dimensional (3D) organoid models have helped overcome the limitations of 2D cell culture, as they display more mature phenotypes, better reflect cell morphology, and recapitulate cell-cell communication networks.^10^

However, it remains difficult to screen drug toxicity in 3D cell culture due to current systems being optimized for 2D monolayered systems. For example, patch-clamp technique relies on isolation of individual cells, which can be difficult to obtain from 3D spheroidal cell aggregates. Additionally, microelectrode arrays assess cells lying on the electrode, and thus are well-suited for 2D monolayers of cells, but contact with 3D spheroidal cell aggregates may be minimal. Furthermore, these methods can be complex, expensive, and resource-intensive with variable results. As an alternative technique, optical analysis of calcium transients, which correspond to action potentials, is generally accepted as an additional method for electromechanical assessment. However, 3D spheroidal organoids may still pose challenges in analysis due to optical signal interference from overlapping cells.

Therefore, we sought to design a novel 2D/3D hybrid organoid system relying on optical techniques and spatial segmentation to obtain electromechanical information on calcium dynamics, cardiomyocyte function, and signal propagation in a high-throughput manner that retains the complexity of a 3D organoid, while also allowing for isolation of single cells. We validated the efficacy of these methods by using a doxorubicin (Dox) treatment model to study the drug’s well-known cardiotoxic, pro-arrhythmic effects.^11^ Furthermore, we utilized machine learning methods to facilitate analysis of the data extracted from our model.

## Results

### hiPSC-CMs demonstrate cardiotoxic effects of doxorubicin

The cardiotoxic effects of Dox on hiPSC-CMs have been well-established, particularly in regard to abnormal calcium handling.^11^ First, we reproduced these findings to confirm that Dox had accurately affected the treatment group. We used the calcium marker Rhod-2 and recorded fluorescence in our model to analyze calcium dynamics (Figure 1a-b). Our optical analysis of calcium transient images revealed that Dox-treated cells had reduced fluorescence peak maximum rate of rise (0.090 ± 0.004 a.u./ms vs. 0.060 ± 0.003 a.u./ms, p<0.0001) and prolonged peak duration (556.2 ± 6.72 ms vs 1063 ± 17.9 ms, p<0.0001), with longer times for both calcium release (182.1 ± 2.74 ms vs 393.6 ± 6.13 ms, p<0.0001) and calcium reuptake (374.1 ± 6.19 ms vs 669.6 ± 16.22 ms, p<0.0001) compared to controls (Figure 1c-f), as has been previously reported.^12,13^ Dox-treated cells also had a greater area under the curve, corresponding to higher levels of cytosolic calcium, compared to controls (2823 ± 119.2 a.u.*ms vs 4745 ± 212.9 a.u.*ms, p<0.0001), which is also consistent with previous reports of Dox-induced calcium overload (Figure 1g).^14^ In alignment with Dox’s pro-apoptotic effects on viability,^15^ the Dox group had decreased spread of cells demonstrated by significantly fewer total surrounding cells per organoid (111.2 ± 6.92 vs 81.29 ± 11.57, p=0.0350, Figure 1h).

**Figure 1:**
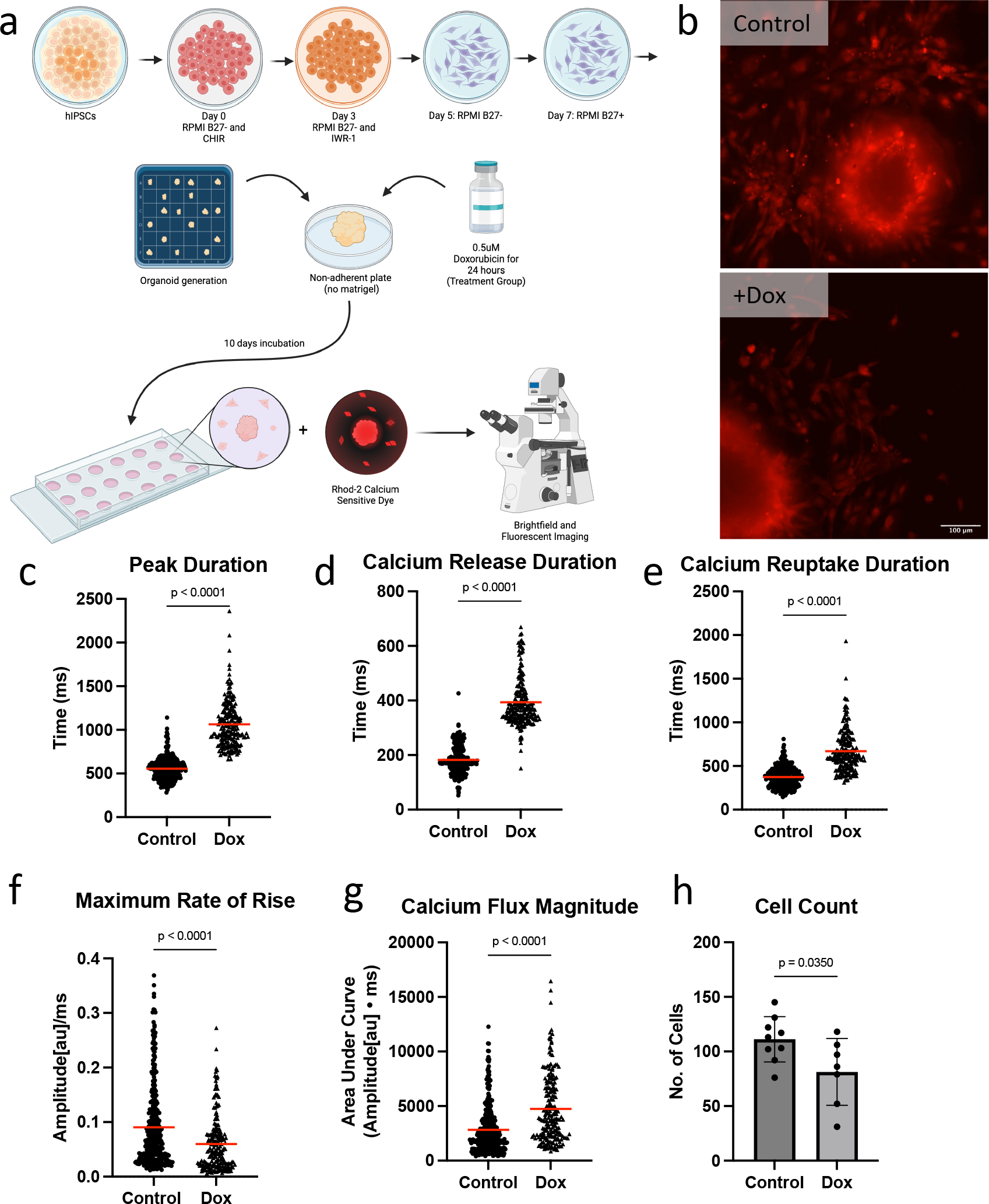
Dox induces detrimental effects on calcium transients in iPSCs. (a) Schematic of experiment. Created with BioRender.com. (b) Representative videos of control and dox organoids and cells with calcium-sensitive Rhod-2 dye. (c) Calcium transient peak duration is prolonged in dox group. (d) Calcium release duration is prolonged in dox group. (e) Calcium reuptake duration is prolonged in dox group. (f) Maximum rate of rise for calcium transient peak is lower in dox group. (g) Area under curve for calcium transient peak, which is indicative of magnitude of calcium, is greater in dox group. (h) The number of cells around the organoid is decreased in dox group.

### 2D/3D hybrid organoid model displays signal propagation effects

To assess calcium signal propagation originating from the organoid, we segmented the area surrounding the organoid and analyzed the same parameters that were used to evaluate overall Dox cardiotoxicity within each region (Center=Organoid, Close=1^st^ adjacent, Mid/Intermediate=2^nd^ adjacent, and Far=remaining outlying region, Figure 2a). Again, the Dox group had decreased propagation of cells, with reduced cellular concentration within each region further from the organoid (Figure 2b). In general, 2D Dox-treated cells had wider waveforms, with diminished peaks with gradual slopes compared to controls and these differences amplified as distance from the organoid increased (Figure 2c). Dox-treated cells had significant prolongation of calcium transient duration as the distance from the center increased (Center vs Close: p = 0.0036, Close vs Mid: p=0.0001), while control cells displayed an increased peak duration only in the farthest region (p<0.0001) (Figure 2d). Similarly, the duration of calcium uptake increased in a stepwise manner in Dox-treated cells (Center vs Close: p=0.0025, Close vs Mid: p<0.0001). Calcium uptake duration also increased in control cells but was only significant between Center and Close (p=0.0462) and Mid compared to Far (p<0.0001) (Figure 2f). In both groups, duration of calcium release did not change across regions (Figure 2e), but maximum rate of rise for calcium release was affected. It was significantly reduced between regions for Dox-treated cells (Center vs Close: p<0.0001, Close vs Mid: p=0.0258), while in control cells, the only significant decrease across adjacent regions was between the organoid and the close region (p<0.0001) (Figure 2g). Spatial alterations of these parameters (Calcium peak duration, release duration, reuptake duration, and maximum rate of rise) on a per-cell basis can be visualized in Figures 2h-k.

**Figure 2:**
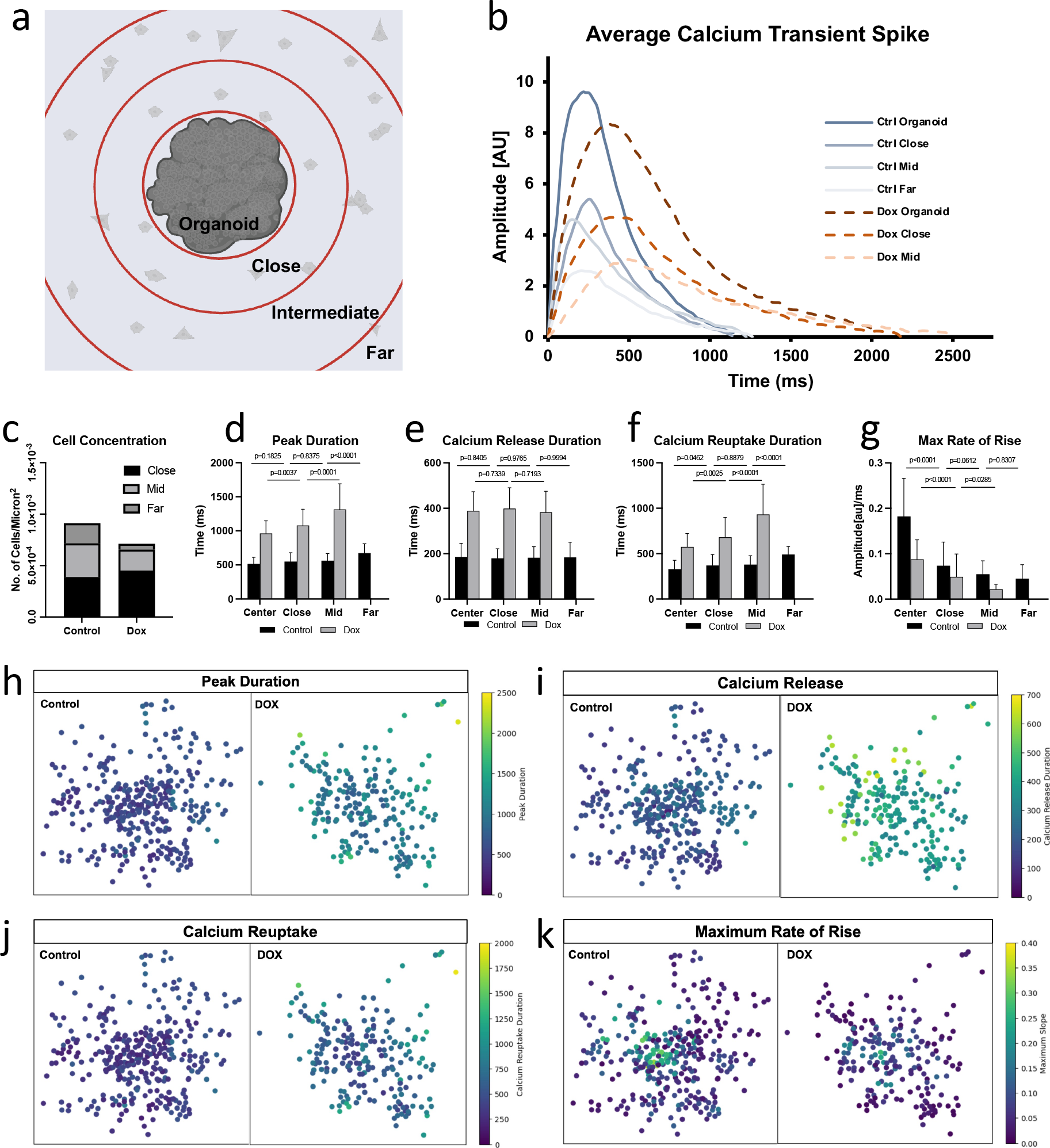
Dox cardiotoxic effects are amplified with increased distance from signal origin (organoid). (a) Schematic of spatial segmentation to assess regional differences. (b) Representative calcium transient peak waveforms for each group. (c) Cellular concentration decreased in the further regions in dox group. (d) Peak duration prolonged with increased distance from the organoid in dox group. (e) Calcium release duration was stable across regions in both groups. (f) Calcium reuptake duration prolonged with increased distance from the organoid in dox group. (g) Maximum rate of rise decreased with increased distance from organoid in dox group. In control group, it only decreased from organoid to first adjacent “Close” region. (h-k) Heat maps of parameters analyzed in d-g. Each dot represents an individual ROI/cell that was analyzed. The center on the graphs was normalized to the center point on each organoid.

### Machine learning to identify abnormal calcium transients

A variety of abnormal calcium transients were observed in both control and Dox-treated cells (Figure 3a), which requires considerable time and effort for thorough examination. Expert classification yielded a total of 639 abnormal cells (58.4%), 291 within controls (49.6%), and 348 within Dox-treated cells (68.6%). Additionally, the proportion of abnormal cells increased within each spatial region further from the organoid in both groups (Figure 3b). We were able to train and test a support vector machine (SVM) model, a widely utilized supervised machine learning model for classification, to classify calcium transients to facilitate burden of this step in analysis (Figure 3c). We used a radial basis function for the kernel type, and the model performed best when tuning the C hyperparameter to a value of 1000. We evaluated the performance of our machine learning classifier based on the confusion matrix, which depicts the number of correctly and incorrectly classified cells (Figure 3d). This allowed us to calculate values for sensitivity (correct abnormal classification), specificity (correct normal classification), and accuracy (total correct classifications), which were 94.7%, 89.7%, and 92.6%, respectively. We then plotted the sensitivity (true positive rate) against the false positive rate (1-specificity) at different classification thresholds to create a receiver operator characteristics (ROC) curve to further demonstrate the accuracy of our model’s classification predictions. The area under the ROC curve value was 0.98, which indicates exceptional performance (Figure 3e).

**Figure 3:**
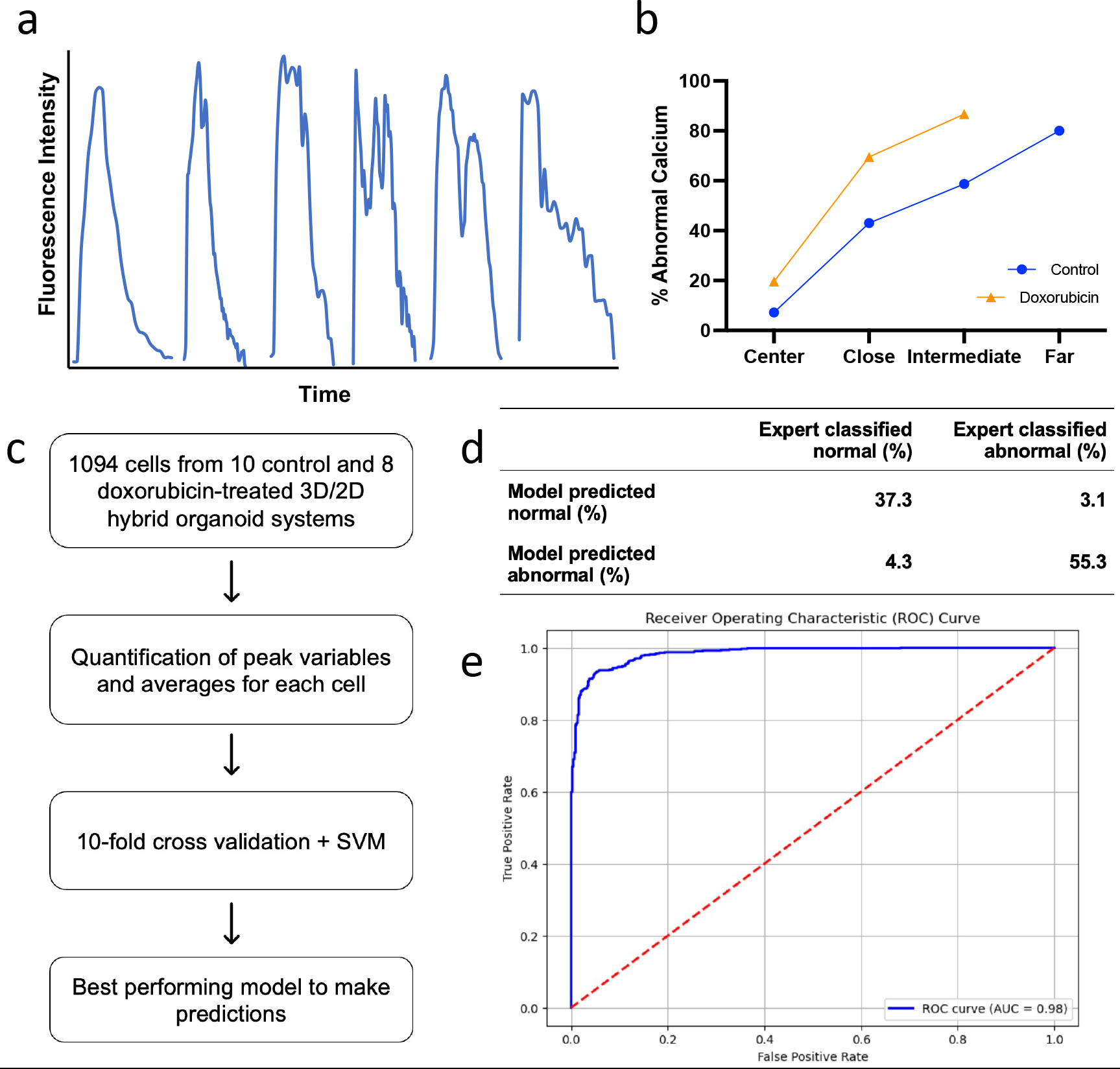
Machine learning accurately classifies abnormal calcium dynamics. (a) The first waveform is an example of normal calcium transient morphology. The remaining waveforms are examples of abnormal calcium transient morphology that was observed. (b) Both control and dox group had abnormal calcium transients and the proportion of cells with abnormal calcium transients increased with further distance from the organoid. Overall, dox had a greater proportion of abnormal calcium transients. (c) Workflow for machine learning classifier model development. (d) Confusion matrix results from best performing model. (e) ROC curve for best performing model, AUC=0.98.

### Dysfunctional contractility reflects Dox-related calcium abnormalities

To assess the effects of Dox and Dox-induced calcium overload on cellular function, we analyzed brightfield images of our model and found that defects in contractility properties correlated with observed calcium abnormalities. First, we found that Dox caused decreased cardiomyocyte function, as a lower proportion of cells per organoid had contractile motion (72.9 ± 5.24% vs 48.5 ± 6.93%, p=0.0125), and there were very few beating cells in the middle and far regions indicating that this effect amplified with increased distance from the organoid (Figure 4b). We also found that among beating cells, 26.4% had dysfunctional contractility in the Dox group, compared to only 15.5% in the control group (Figure 4c). To assess the correlation between dysfunctional contractility and calcium dynamics, we analyzed the calcium transient data for the cells with abnormal beating patterns and found that both groups had similar proportions of abnormal calcium transients (71.9% in control vs 74.8% in Dox group, Figure 4c). Total beat, systole, and diastole durations were significantly prolonged to a similar extent in our Dox-treated cells (total beat duration: 0.410 ± 0.005 s vs 0.688 ± 0.012 s, p<0.0001; systole: 0.159 ± 0.003 s vs 0.293 ± 0.011 s, p<0.0001; diastole: 0.252 ± 0.004 s vs 0.400 ± 0.009 s, p<0.0001; Figure 4d-f). In the control group, durations for systole and diastole varied across the different spatial regions, but there were no consistent findings of prolongation or shortening based on distance from the organoid. Conversely, we observed that total beat time and systole were longer in the surrounding cells than organoid in our Dox-treated cells (p<0.05). Correlating with prolonged calcium reuptake, we also observed a consistent extension in diastole duration, although this did not reach statistical significance (Figure 4g-l).

**Figure 4:**
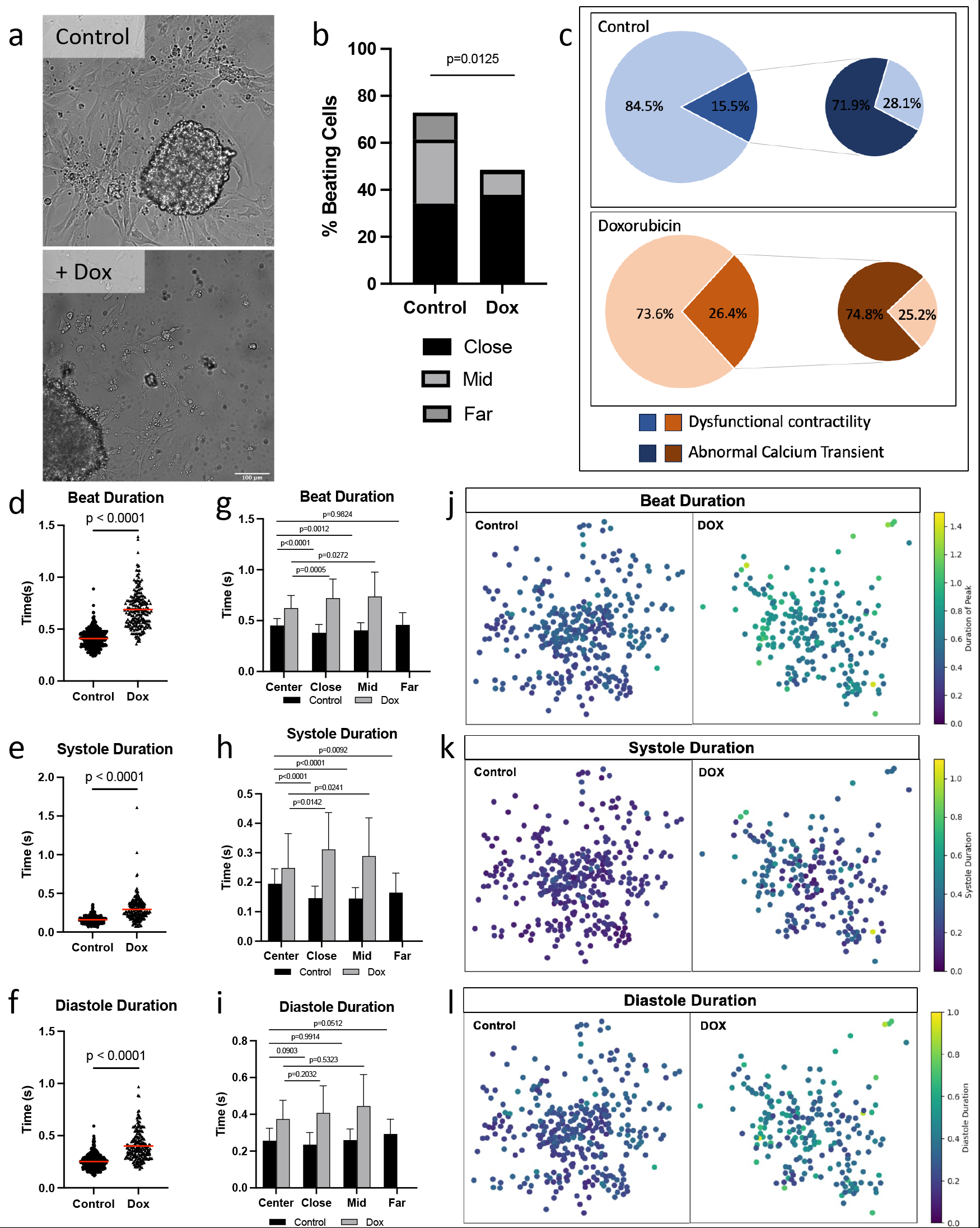
Dox-related calcium abnormalities are mirrored in CM contractility. (a) Brightfield videos of control and dox organoids and cells. (b) Dox group had fewer beating cells, which were mostly concentrated in close region. (c) Dox had more cells with dysfunctional contractility, but in both groups a similar percentage of dysfunctional beating can be attributed to abnormal calcium handling. (d) Total beat duration is prolonged in dox group. (e) Dox group has prolonged systole. (f) Dox group has prolonged diastole. (g-i) Beat, systole, and diastole durations have temporospatial changes. (j-l) Heat map representation of temporospatial changes on a per-cell basis.

## Discussion

Due to the need for improved assessment and prediction of cardiotoxicity during drug development and testing, we have developed a novel 2D/3D hybrid organoid model which uses optical imaging techniques to study drug effects via calcium transients, contractility, and, most distinctly, signal propagation. We found that our novel 2D/3D hybrid organoid model accurately assessed the known cardiotoxic effects of Dox, and thus, presents a feasible method for evaluating drug cardiotoxicity on a cellular level in a high-throughput manner. To first validate our model, we assessed Dox cardiotoxicity and confirmed its known effects. Calcium transient data demonstrated prolonged total peak durations, extended release and reuptake phases, reduced frequency, and decreased maximum rate of rise. These results align with case studies of patients arrythmias, such as QT interval prolongation, extrasystoles, atrial fibrillation, and ventricular arrythmias, all of which can in part be attributed to defects in calcium handling.^16-19^ Previous studies in animal models and in 2D and 3D iPSC cultures have also demonstrated decreased heart rate, prolonged QT interval, and irregular beating pattern, as well as altered calcium dynamics, including decreased reuptake and increased concentration, providing further validation to our results using calcium transients as a surrogate marker of cardiotoxicity.^12-14,20^

Each of these previously used models, however, have drawbacks, namely related to consistency and feasibility. Our novel 2D/3D hybrid organoid model provides a more robust and optimal platform by optimally balancing the advantages and disadvantages of each. For example, results found in 2D cell culture have yielded contradictory effects, with some studies finding reduced beating rate and prolonged action potential duration, while others have reported increased frequency, no acute effects on beating, or a complete cessation of contractile motion.^14,20,21^ Additionally, studies have shown that 2D cultured cells lack the complexity seen in real tissues and have divergent transcriptomes and electromechanical properties, which offers explanation to different toxicity responses observed in 2D versus 3D cell culture.^9,10,22-25^ Our model circumvents these discrepancies because the isolated single “2D” cells in our hybrid model are developed as part of the organoid, and not as monolayers, thus preserving properties of true cardiomyocytes and cell-cell interactions, while still allowing for easy optical analysis without interfering adjacent signal. Overall, our hybrid model overcomes the challenges of other systems by presenting reliable, easy-to-interpret results in human cells that display the complexity and structure of cardiomyocytes.

Furthermore, what distinguishes our hybrid model from others is the dispersion of cells around the beating organoid, which allows us to study signal propagation and how the signal changes from its origin on a per cell basis. This is important because a major contributor to arrhythmias is conduction velocity, which is hypothesized to be impaired in Dox cardiotoxicity.^11,26^ Our results demonstrated diastole prolongation with increased distance from the beating organoid, suggesting that Dox cardiotoxicity amplifies across a signal. This can contribute to arrhythmias due to calcium overload impacting the initiation of the next action potential, which can predispose cells to altered signals and arrhythmias. Correlative findings have also been previously reported in human patients, 2D and 3D cell culture, and in animal models, which have shown increased QT dispersion, increased field potential duration, increased dispersion of repolarization, and reduced and heterogenous conduction velocity.^14,27-31^ Although these findings report different metrics, they converge on the underlying observation that signal propagation is not uniform throughout, and such signal dispersion and electrical heterogeneity with nonuniform repolarization and refractoriness can induce arrhythmias. Mechanistic investigations have supported these conclusions, demonstrating Dox and calcium overload impact connexin-43, the main gap junction protein found in cardiac tissue.^32-35^ Specifically, Dox has been found to impair expression and function of connexin-43, which can lead to conduction velocity slowing and heterogeneity, while calcium overload has been shown to exacerbate conduction slowing through down-regulated gap junction communication and electrical uncoupling.^33-39^ Thus, there are several mechanisms that contribute to cell-cell communication and offer explanation to the impaired signal propagation demonstrated in our model.

Additionally, our results reflect what has been elucidated from molecular studies of Dox cardiotoxicity and abnormal calcium handling. Firstly, Dox is known to cause a pro-apoptotic environment and decrease cell viability, which explains why we have overall fewer cells in our Dox-treated models.^14,15,34^ The prolonged time for calcium reuptake and overall increased levels of calcium found in our study are reflective of previous work identifying dysfunctional SERCA2a proteins as a cause for impaired calcium release and reuptake in Dox cardiotoxicity.^12,35,40,41^ In addition to direct effects seen on SERCA2a expression, Dox has also been shown to increase reactive oxygen species, which are known to alter SERCA2a activity and cause diastolic calcium leak causing contractile dysfunction and impaired cardiomyocyte relaxation, which we have also observed in our model.^15,42,43^ Impaired SERCA2a expression and function is a proposed mechanism for abnormal contractility and has also been seen in heart failure, arrhythmias, and ischemic heart disease, and therefore is a likely contributor to Dox cardiotoxicity.^44-46^

Considering the tight control of excitation-contraction coupling, it is not surprising that our Dox treated cells also display impaired contractility that is reflective of the effects seen on calcium transients. Previous studies have elucidated that calcium transients decay rate closely mirrors myocyte contraction, implicating that decreased calcium reuptake and subsequent elevated intracellular calcium levels during diastole play a significant role in impaired and delayed relaxation.^47^ Specifically in Dox models, depressed contractility and impaired relaxation have been attributed to altered sarcoplasmic reticulum calcium release and uptake.^48,49^ Furthermore, Dox and its associated calcium overload have been shown to reduce contractile structures and impair their function due to sarcomeric disarray and myofibril deterioration.^50-53^ This provides explanation for why some of our cells with dysfunctional contractility did not have calcium abnormalities. On the other hand, studies have shown that calcium transients become abnormal significantly earlier than contractility defects arise, explaining why not all cells with abnormal calcium had abnormal beating.^54^ Therefore, our model provides additional insight into excitation-contraction coupling impairment in Dox cardiotoxicity and can clarify if its attributable to calcium, cellular architecture, or both.

Given that many of the detrimental effects caused by Dox can be attributed to abnormal calcium handling, this study emphasizes the need to correctly detect normal versus abnormal calcium transients. When done manually, this process is burdensome, inefficient, and prone to human error due to borderline signal, output noise, subjectivity, and expertise. Therefore, we additionally developed a machine learning classifier for assessing normality. Previous studies have shown the ability of various machine learning methods, including SVM, to successfully classify cells based on calcium transient data. This has been done to assess impaired relaxation, predict mechanistic action of cardioactive drugs, classify different cardiac diseases, and categorize normal versus abnormal calcium transients.^55-58^ These models have all had impressive accuracy ranging from 80-88%, and our model’s had comparable results with an accuracy improved to 92.6%. Because irregularities may relate to not solely to transient morphology but also to signal phase and because cells may occasionally have a singular abnormal signal amongst normal transients and vice-versa, we classified calcium transients on a whole cell basis, based on averaged parameters, rather than a single peak basis to obtain the most accurate classification for a cell to assess overall cardiotoxic burden. Additionally, since both control and Dox-treated cells had calcium transients of normal and abnormal morphology, we trained and tested our model to classify calcium transients regardless of drug-treatment status to capture underlying patterns of calcium abnormality in an unbiased manner to ensure it was not solely classifying based on drug-status. This improves its generalizability among cardiomyocytes with different cardiotoxicities, which will enhance its ability to screen a variety of drugs in a high-throughput manner.

Despite overcoming the challenges of previously utilized model systems for assessing drug cardiotoxicity, our study is subject to the following limitations. We assessed one drug at one concentration at one timepoint. Future studies employing our model with numerous drugs at a variety of concentrations at different time points will provide further validation and proof of utility. Given its ability to screen drugs in a high-throughput manner, this can easily be achieved. Additionally, we imaged calcium transients and contractility consecutively as opposed to simultaneously. Assessment of these two parameters in conjunction can be done with this model with the proper equipment and would shed further light on a drug’s impact on excitation-contraction coupling.

In conclusion, our 2D/3D hybrid organoid model has potential to offer a novel, high throughput, low cost, and effective drug screening tool that can be utilized to assess several factors impacting cardiac health, such as calcium transients, contractility, and signal propagation. Given that most cardiotoxic effects relate to pro-arrhythmia, our model is well-suited to study most drugs. It additionally allows for studying contractility in association with electrical effects. The isolation of single cells can also allow for staining and assessment of other structures impacted in drug-related cardiotoxicity such as mitochondria, cellular architecture, and alpha actinin. Furthermore, it has potential to provide personalized, precision medicine by utilizing iPSCs directly from patients to study how they may uniquely react to potential drug candidates.

## Methods

### Cell culture and model development

Wild type hiPSCs were obtained from NIH iPSC repository (ND2.0, NIH CRM control iPSC line) and plated on Matrigel coated plates until they reached 80-90% confluence at which point they were split and transferred to a 6 well plate. iPSCs were differentiated using 2mL RPMI B27 minus and 6uM CHIR for 2 days. At 96 hours media was replaced with RPMI B27 minus and 5uM IWR-1. After 48 hours, they received RPMI B27 minus, which was replaced by RPMI B27 plus after an additional 48 hours. At day 7, cells were transferred to an AggreWell plate to generate organoids. After cells were added to the AggreWell plate, the plate was centrifuged and incubated at 37 degrees Celsius for 48 hours. Subsequently, organoids were dislodged from the AggreWell plate and transferred into non-adherent plates and maintained in RPMI B27 media. Organoids were treated with 0.5uM Dox or control media for 24 hours and then switched to maintenance media with RPMI B27 for 10 days following drug treatment. At that point organoids were placed in ibidi wells with Matrigel to allow for single cells to dissociate from the organoid for further evaluation. The cells were then stained with Rhod-2 for calcium analysis. A 2.5mM stock of Rhod-2 was diluted 1:1000 in RPMI B27 plus, added to cells, and incubated at 37 degrees Celsius for 15-25 minutes after which they were washed two times with RPMI B27 plus.

### Image Capture and Video Analysis

Fifteen-second videos of unpaced, spontaneously beating cardiomyocyte organoids were captured at 20x using confocal microscopy with absorbance and emission fluorescent staining. To visualize calcium transients within cells, calcium indicator Rhod-2 was used, and then cells were stained with Cy3. Fluorophores were excited at and imaged at appropriate emission wavelengths (590nm, red). Brightfield videos were consecutively recorded also at 20x for fifteen seconds. Videos were visually assessed for adequate distribution of 2D cells around the organoid. Videos that lacked surrounding cells or fluorescence were excluded from analysis for a total of eighteen (10 control, 8 Dox) videos analyzed. Videos were uploaded into ImageJ, and the “ROI Manager” tool was utilized to select contracting cells within each brightfield video and discrete areas of Rhod-2 excitation/emission within each fluorescent video, with an average of approximately 65 ROIs/cells identified per video.

### Calcium Analysis

To assess calcium transients, we utilized an Image J plugin called Spiky, which is a software tool that quantifies excitation–contraction related experimental data, including intracellular calcium transients.^59^ It detects peaks in the input by characterizing the temporal shift from baseline of a signal in an image, and then computes a variety of parameters for each peak, such as amplitude, time to peak, time between peaks, peak width, maximum slope of peak, and area under the curve. Spiky was run for each individual ROI. Default settings were used for peak detection. Full width and right and left half widths at 20% maximum represented total flux, efflux, and influx. Area under the curve is a function of fluorescence amplitude and time and thus is representative of relative calcium signal magnitude. Drift removal was applied to all outputs to normalize baseline shifts. Graphical results were evaluated for quality control (correct detection of calcium dynamic peaks).

### Contractility Analysis

To further evaluate our model’s reliability, we analyzed brightfield videos to assess contractility, which should mirror the effects seen in calcium transient analysis based on excitation-contraction coupling. To analyze contractility, we utilized an Image J macro called Myocyter.^60^ This tool characterizes contracting regions using two parameters: “speed,” which is the difference between consecutive frames of a video and indicates the speed of a contraction, and “amplitude,” which is the difference between the current frame and an automatically determined reference frame of the cell in its resting phase and indicates the amount of deformation of the contracting cell compared to the reference frame. The difference between the images is then calculated and presented as a plot and numerical output, generating values for mean frequency, amplitude, systole (contraction time), diastole (relaxation time), peak times (total length of contraction and relaxation), and beat time (peak to peak time). Using dynamic thresholding, the program precisely tracks amplitudes even when the baseline shifts. Myocyter was run using the manual ROI list and default settings for intensity threshold, particle size, peak recognition, and sensitivity of maxima and minima detection. Analysis was performed on a per-cell basis, and graphical results were evaluated for quality control (correct detection of contractile motion).

### Regional/Spatial Segmentation

To analyze spatial differences in calcium transient and contractility parameters based on distance of 2D surrounding cells from the beating organoid, we segmented images into regions of equally increasing distance from the beating organoid. Three concentric circles centered on the organoid were overlaid on each frame, each with a distance of 185 microns between. The regions were labeled “Center” for the organoid, “Close” for the first adjacent region, “Mid/Intermediate” for the second region from center, and “Far” represents the remaining area in the frame of the image (Figure 2a).

### Machine Learning

First, evaluation of calcium transient abnormality was conducted by two experts for the signal from each cell. In instances of discordance among the two experts, a third expert inspected the signal to resolve discrepancies. A cell was labeled as having abnormal calcium transients if it had substantial atypical calcium transient morphology, such as bifid or multi-spiked peaks, plateauing of calcium signal during uptake, increasing calcium during uptake as indicated by a second upstroke occurring prior to complete signal decay, and evidence of spontaneous calcium release between peaks. Next parameters obtained from Spiky were used to train the SVM. To ensure our model was not overfitting data, we used 10-fold cross-validation. Additionally, we tuned a regularization hyperparameters of the classifier model, denoted by *C*, using a grid search within a range of 0.001-1000. To assess the performance of the model, we relied on sensitivity, specificity, and accuracy calculations, as well as the area under the ROC curve.

### Statistical analysis

Statistical analyses were performed in GraphPad Prism 10.2.1. All data are reported as mean ± standard error of the mean unless noted otherwise. Two-tailed Student’s t-tests were performed to compare control and Dox-treated groups. One-way ANOVA followed by Tukey’s post-hoc test was used to detect differences across spatial regions. A p-value of less than 0.05 was considered statistically significant.

## Data availability

The data reported in this study are available and can be requested by contacting the corresponding author.

## Author Contributions

JL: Conceptualization, Software, Formal analysis, Data curation, Writing - Original Draft, Visualization.

BY: Conceptualization, Methodology, Investigation, Writing - Review & Editing.

AS: Conceptualization, Methodology, Resources, Writing - Reviewing & Editing, Supervision, Project administration.

Each author contributed intellectually to the design of the study and interpretation of data. All authors have reviewed and approved the manuscript.

